# Small activating RNA AW1-51 (CEBPA-51) elicits targeted DNA demethylation to promote gene activation

**DOI:** 10.1101/2025.09.29.679274

**Authors:** Giulia Gaggi, Juan Rodrigo Patiño-Mercau, Marta Borchiellini, Mailin Li, Lucrezia Rinaldi, Giorgia Maroni, Davide D’Onghia, Susumu S. Kobayashi, Mahmoud A Bassal, Nagy A. Habib, Pedro P Medina, Angela Di Baldassarre, Barbara Ghinassi, Alexander K Ebralidze, Simone Ummarino, Daniel G Tenen, Annalisa Di Ruscio

**Affiliations:** Harvard Medical School Initiative for RNA Medicine, Harvard Medical School, Boston, MA, 02115, USA; Cancer Research Institute, Department of Medicine, Beth Israel Deaconess Medical Center, Harvard Medical School, Boston, MA, USA; Department of Medicine and Aging Sciences, “G. d’Annunzio” University of Chieti-Pescara, 66100 Chieti, Italy; Houston Methodist Neal Cancer Center, Houston Methodist Hospital, Houston, TX, USA; Texas A&M MD/PhD Program, Texas A&M College of Medicine, Bryan, TX, USA; Institute of Biomedical Technologies, National Research Council (CNR), Pisa, Italy; Department of Translational Medicine, Università del Piemonte Orientale (UPO), 28100 Novara, Italy; Department of Surgery & Cancer, Imperial College London, W12 0NN London, UK; MiNA Therapeutics Ltd, W12 0BZ London, UK; Gene Expression Regulation and Cancer Group (CTS-993), GENYO, Centre for Genomics and Oncological Research: Pfizer-University of Granada-Andalusian Regional Government, Granada, Spain; Department of Biochemistry and Molecular Biology I, Facultad de Ciencias, University of Granada, Avda. de Fuentenueva S/N, 18071, Granada, Spain; Health Research Institute of Granada (Ibs.Granada), Granada, Spain; Reprogramming and Cell Differentiation Lab, Center for Advanced Studies and Technology (CAST), 66100 Chieti, Italy; Department of Medicine and Aging Sciences, University “G. d’Annunzio” of Chieti-Pescara, 66100 Chieti, Italy; Department of Innovative Technologies in Medicine and Dentistry, “G. d’Annunzio” University Chieti-Pescara, Chieti, Italy; Harvard Stem Cell Institute, Harvard Medical School, Boston, MA, USA; Department of Biology, Tufts University, Medford, MA 02155, USA

**Author notes:** These authors contributed equally to this work (Giulia Gaggi, Juan Rodrigo Patiño-Mercau). These authors contributed equally to this work (Marta Borchiellini, Mailin Li).

**Keywords:** Small activating RNA, DNA methylation, *CEBPA*, cancer, AW1-51

## Abstract

Small activating RNAs are short double-stranded RNAs designed to upregulate transcription of target genes. By this virtue, they can be used to restore expression of genes frequently silenced in cancer. AW1-51 (also referred to as CEBPA-51), the first small activating RNA therapeutic to enter clinical evaluation, has demonstrated biological activity and safety in Phase II trials for hepatocellular carcinoma, both as monotherapy and in combination with sorafenib, and in Phase 1a/1b in combination with pembrolizumab for patients with advanced solid tumors. It targets the master regulator CCAAT enhancer-binding protein alpha, abnormally silenced by DNA methylation in a wide range of hematological and non-hematological malignancies. However, the molecular events enabling this mechanism are only partially elucidated.

In this study, we uncovered the molecular basis for AW1-51-induced transcriptional reactivation of CCAAT enhancer-binding protein alpha demonstrating that by directly promoting DNA demethylation of its promoter restores its expression, protein synthesis, and consequently cell differentiation.

These findings unveil AW1-51 as a prototype for RNA-based precision medicine enabling conditional expression of CCAAT enhancer-binding protein alpha in diseases characterized by aberrant gene silencing and extending its potential therapeutic impact beyond cancer.

## Introduction

Small activating RNAs (saRNAs) represent a novel class of synthetic double-stranded RNAs capable of upregulating gene expression by targeting specific promoters i.e., *E-cadherin* (1), *P21* (2, 3), *Progesterone Receptor* (4), pluripotency genes (5–7).

AW1-51 (also referred to as CEBPA-51) was the first saRNA therapeutic to enter clinical evaluation and demonstrating biological activity and safety in Phase I and II clinical trials (ClinicalTrials.gov: NCT02716012; EudraCT 2021-005431-23, respectively) for hepatocellular carcinoma, both as monotherapy and in combination with sorafenib (8, 9). Also, AW1-51 is currently being evaluated in a Phase 1a/1b clinical trial in combination with pembrolizumab (a PD-1 inhibitor) for patients with advanced solid tumors (ClinicalTrials.gov: NCT04105335) (10–12).

AW1-51 targets the coding sequence of CCAAT/enhancer-binding protein alpha (*CEBPA*), and it is, so far, the only reported saRNA that does not target the gene promoter (Figure 1a). *CEBPA* is a methylation-sensitive gene that encodes a key transcription factor involved in myeloid lineage development and liver function. *CEBPA* expression is often silenced by DNA methylation in leukemias, liver, and lung cancer making this *locus* therapeutically appealing (13–15).

**Figure 1.**
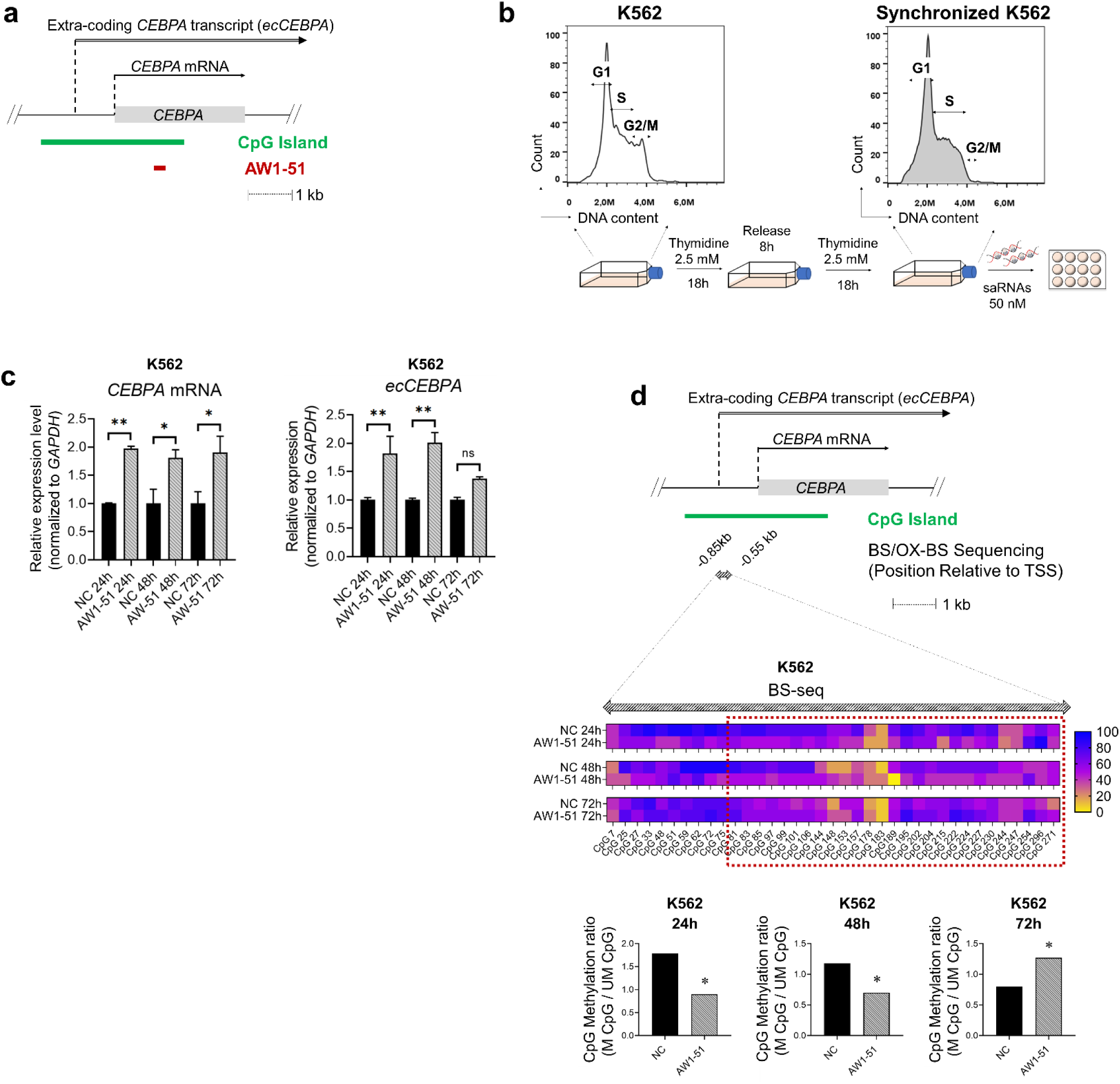
AW1-51 induces gene expression and promoter demethylation in a silenced CEBPA locus. **a,** Schematic of the human CEBPA locus and transcripts. Green rectangle indicates the CEBPA associated CpG island. Red rectangle indicates the AW1-51 target sequence. **b,** Schematic of saRNA delivery into K562 cells. K562 cells are arrested at the G1/S phase boundary by double-thymidine block (2.5 mM; 18h block, 8h release, 18h block). After synchronization, cells are electroporated with 50 nM of AW1-51 or a nontargeting control saRNA (NC). c, RT-qPCR results showing CEBPA (left) and ecCEBPA (right) expression from 24h to 72h post transfection in K562 cells (n=2). One-way ANOVA. *: p<0.05, **: p<0.01, ns: non-significant. d, Region within the CEBPA CpG island (striped double-headed arrow, -0.85/-0.55kb before CEBPA’s transcription start site [TSS]) interrogated by bisulfite sequencing (BS-seq) in K562 cells. Changes in DNA methylation are shown as heatmaps (see detailed BS-seq results in Supplementary Figure 1d). Bar graphs show the ratio between methylated (M) or unmethylated (UM) CpGs to total CpGs within the region included in the dashed line red box. All bisulfite sequenced clones were analyzed by Fisher’s exact test (n=12). *: p<0.05.

DNA methylation and demethylation are dynamic processes that play crucial roles in gene regulation. The conversion of cytosine to 5-methylcytosine (5mC) typically leads to transcriptional silencing and is catalyzed by enzymes of the DNA methyltransferase family (DNMTs). This methylation process is commonly found at CpG dinucleotides, particularly in regions known as CpG islands (CGIs), located near the transcriptional start site (TSS) of most protein-coding genes in humans. In contrast, the removal of 5mC to activate gene expression involves the oxidation of 5mC to 5-hydroxymethylcytosine (5hmC), a reaction carried out by the Ten-Eleven Translocation (TET) family of dioxygenases. The conversion of 5mC to 5hmC is a key intermediate step of the active demethylation process and is often associated with the activation of previously silenced genes (16, 17). Alterations of the pathways regulating the balance between DNA methylation and hydroxymethylation patterns have been implicated in various human diseases including cancer, and are currently under intense investigation as potential therapeutic targets in both solid and liquid tumors (18, 19).

In this instance, RNAs interacting with epigenetic writers, such as DNA methyltransferases (DNMT1), or erasers i.e., Ten-eleven translocation (TET) family proteins were shown to regulate gene expression and are considered modulators of epigenetic machinery (20–23).

Despite the evidence showing saRNAs to regulate gene expression, their role in modulating epigenetic pathways, particularly through DNA methylation and demethylation, has never been fully investigated. saRNAs are thought to act at various levels by: promoting chromatin accessibility via histone modifications (4), releasing promoter-proximal pausing (3), and/or titrating the balance between sense and antisense transcripts (2, 5). However, whether saRNAs like AW1-51 control transcriptional regulation through participation at the epigenetic layer remain largely unclear.

Herein, we hypothesize that AW1-51 restore *CEBPA* transcription by eliciting targeted DNA demethylation the *CEBPA locus* and promoting gene activation.

More broadly, our findings might suggest a new layer of control for AW1-51 and its potential application in diseases driven by aberrant DNA methylation patterns.

## Materials and Methods

### Cell culture

K562, A549, U937 and HL60 cell lines were grown in RPMI medium supplemented with 10% fetal bovine serum (FBS; Gibco, Thermo Fisher Scientific). All cell lines were purchased from ATCC and grown at 37°C (21% O2, 5% CO_2_) in the absence of antibiotics.

### 5-Azacytydine treatment

For K562, 400,000 cell/mL were seeded in RPMI 10% FBS medium supplemented with 10μM of 5-Azacitidine (5-Aza) (Sigma-Aldrich). For A549, 50,000 cell/mL were seeded in RPMI 10% FBS and incubated overnight to allow cell adhesion. The next day, 15μM 5-Aza was added to the cell culture. Fresh medium with 5-Aza was replaced for both K562 and A549 after 48h. DNA and RNA were collected for downstream approaches after 72h for K562 and 96h for A549.

### DNA Isolation

Genomic DNA was isolated using a homemade Lysis buffer (10 mM Tris pH 8, 25 mM EDTA pH 8, 0.5% SDS, 200 mM NaCl) followed by RNase A treatment (25μg/mL, Roche) for 30 minutes at 37°C and then Proteinase K (0.44mg/mL, Roche) overnight at 65°C. DNA was then recovered by basic phenol:chloroform:isoamyl alcohol 25:24:1 pH 8.0 (Sigma) extraction and 1:1 isopropanol precipitation. DNA was resuspended in TE pH 8.0 overnight and quantified by Qubit 3 (Thermo Fisher Scientific) following the manufacturer’s procedures for all downstream applications.

### Bisulfite and Oxidative-Bisulfite sequencing and combined bisulfite restriction analysis (COBRA)

Bisulfite sequencing was performed as previously described (21). Briefly, 200 ng to 1 ug of genomic DNA was bisulfite converted for 7.5 hours using the EZ DNA Methylation kit (Zymo Research) and eluted in water. For Oxidative-bisulfite sequencing, 200 ng of DNA was converted by TrueMethyl® oxBS-Seq Module (Tecan) following the manufacturer’s procedure. PCR amplification of bisulfite and oxidative-bisulfite-treated DNA was performed with FastStart Taq DNA Polymerase (Roche). Reaction conditions were 95°C (6 min), followed by 35 cycles at 95°C (30s), 53-57°C (1 min), and 72°C (1 min), with a final elongation at 72°C (7 min). Primer sequences for bisulfite/oxidative bisulfite sequencing and COBRA are summarized in Supplementary Data 1.

For bisulfite and oxidative-bisulfite sequencing, the gel-purified PCR product was cloned into the pGEM-T Easy Vector System (Promega) and screened by blue/white colony screening and antibiotic resistance. Sequencing results were analyzed using QUMA software (http://quma.cdb.riken.jp/) (24). Clones with percentage of unconverted CpGs >95 and with percentage of identity <90 were excluded from the final analysis. The minimum number of clones for each sequencing condition was 12.

For COBRA, 400 ng of 2% TAE agarose gel-purified PCR product (QIAEXII Gel Extraction Kit, Qiagen) was digested with 10 units of BstU1 enzyme (New England Biolabs) at 60°C for 2 hours. Digestion products were analyzed on a 3% TAE agarose gel.

### Early S Phase Cell Cycle Synchronization

Thymidine (Sigma Aldrich) was prepared in cell culture water at 50 mM concentration, sterilized by 0.22-micron (µm) filtration, and then stored at -20°C until use. K562 cells were seeded at 0.5×10^6 cell/ml and the synchronization followed the 18-8-18 h time course: 2.5 mM thymidine – release in complete RPMI - 2.5mM thymidine. For A549, a one-time block of 2 mM thymidine for 18 hours was sufficient to synchronize 2.3×10^6 cells (25).

Both cell lines synchronization was confirmed with Pyronin Y (1.5 mg/ml in PBS, Sigma) and Vybrant DyeCycle Violet (Invitrogen) stained according to manufacturer’s recommendations. Flow cytometry was performed using the CytoFLEX LX (Beckman Coulter) in the B610-ECD filter (488 nm laser) and V450-PB filter (405 nm laser).

### Small activating RNAs (saRNAs) treatment

AW1-51, targeting the coding region of the gene, is the same as previously published (26) and under current clinical investigation (Clinicaltrials.gov: NCT02716012) (9). A non-targeting saRNA was used as negative control (NC) in each transfection. Pre-annealed sequences were ordered from Sigma, resuspended to 100 μM concentration with cell culture grade water, and kept at -20°C until use. saRNA sequences are listed in Supplementary Data 1.

saRNAs (50 nM) were transfected into K562 cells with Kit V program T-016 by Amaxa Nucleofector Device II (Lonza). In A549, saRNAs (50 nM) were transfected using DharmaFECT 1 (DF1, Horizon Discovery). Following Voutila et al protocol, cells were first reverse transfected and then forward transfected 24h later (26). Transfection efficiency was monitored by flow cytometry or fluorescent microscopy based on expression of Green Fluorescence Protein encoding plasmid compared to non-electroporated / non-transfected control. For the experiments that required a larger number of cells, the ratio of cells to saRNA molecules remained consistent throughout.

### RNA extraction

Total RNA was isolated by phenol chloroform purification using the commercially available TriReagent (Ambion, Thermo Fisher Scientific) according to manufacturer’s recommendations. After 1:1 isopropanol precipitation of the aqueous phase, the RNA pellet was treated with DNAse I (Roche; 15 U per sample) at 37° C for one hour in the presence of RNase Inhibitor (Promega). DNase was inactivated with the addition of 10 mM EDTA pH 8 (Sigma-Aldirch) and RNA was then recovered by acidic phenol (Sigma, pH 4.3) extraction and 3:1 ethanol precipitation in the presence of 200 mM NaCl. RNA was resuspended in RNAse-free water and quantified by Qubit 4 for all downstream applications.

### Reverse transcription-quantitative polymerase chain reaction (RT-qPCR)

TaqMan-based RT-qPCR was performed using the iTaq Universal Probes One-Step Kit (Bio-Rad) 100 ng of total RNA. Reaction conditions were 50°C (10 min), 95°C (2 min), followed by 40 cycles of 95°C (15s) and 60°C (1 min) with fluorescence acquisition during the final step. GAPDH or 18s were used as normalization controls (Applied Biosystems; Thermo Fisher Scientific). Target gene amplification was calculated using the formula 2^-ΔΔDDCt as described by Livak et al (27). All primer and probe sequences are listed in Supplementary Data 1.

### Actinomycin D treatment

Synchronized A549 cells were transfected with saRNAs (negative control and AW1-51) as described above, using 130,000 cells per condition and timepoint. After transfection, cells were treated with 1 µg/mL Actinomycin D (Sigma-Aldrich) to inhibit transcription. Total RNA was extracted at 30 min, 1 h, 3 h, 6 h, 8 h, and 24 h post-treatment, and transcript stability was assessed by RT-qPCR.

### Western blot

Single cells suspensions were lysed in Reduced 2x Sample Buffer (0.1 M Tris pH 6.8, 4% SDS, 5% glycerol, 0.2 M DTT and 4% β-mercaptoethanol) according to cell number and then boiled at 95°C for a minimum of 5 minutes. Equivalent amounts of protein were resolved in 12% polyacrylamides gels and transferred to polyvinylidene fluoride (PVDF) membranes (Bio-Rad). Blots were blocked with 5% nonfat dry milk in TBS 0.05% TWEEN 20 (0.05% TBS-T) prior to incubation with primary antibodies. Antibodies were diluted as following: CEBPA (Cell Signaling, #2295S) 1:1000 in 5% BSA in 0.1% TBS-T and β-Actin (Santa-Cruz Biotechnology, #sc-81178) 1:5000 in 5% nonfat dry milk 0.1% TBS-T. Secondary antibodies were diluted as following: anti-rabbit (Bio-Rad, #1706515) 1:5000 and anti-mouse (Abcam, #ab6789) 1:3000. They were visualized by enhanced chemiluminescence detection. Bands quantification was performed using Fiji (ImageJ) software (28). Data was normalized with β-actin values.

### RNA sequencing

Total RNA samples from A549 and K562 cells were collected 72h after transfection with the saRNAs. Two samples per condition were prepared and sent to BGI Genomics Inc. The poly A RNA fraction was extracted and RNA libraries were prepared using Stranded total RNA libraryIllumina Truseq stranded kit / MGI’s directional kits, and sequenced in paired end. Raw fastq files first had optical duplicates removed using clumpify from the BBTools (sourceforge.net/projects/bbmap/) suite using the flag “dedupe spany addcount”. Next, quality trimming was performed using Trimmomatic (version 0.36); Bolger et al., 2014) in paired end mode with the flags “LEADING:20 TRAILING:20 SLIDINGWINDOW:4:15 MINLEN:36”. Finally, reads were filtered according to their overall quality using fastq-filter (https://github.com/LUMC/fastq-filter) with the flag “-e 0.01”, which corresponds to the average error rate of 0.01 per base (phred score of 20). Alignment was performed using STAR (version 2.5.4a; Dobin et al., 2013) with the flags “--outFilterScoreMinOverLread 0.1 -- outFilterMatchNminOverLread 0.1 --outFilterMultimapNmax 2” with the resultant bam files sorted and indexed using samtools. FeatureCounts feature of the Subreads (version 2.0.0; Liao et al., 2014) package was used to generate a count matrix taking into account the paired end read and allowing for multimapping of the reads. Differential expression analysis was performed using the DESeq2 (version 1.44.0; Love et al., 2014) in R and plotted using EnhancedVolcano (version 1.22.0; Blighe et al., 2025) with a significant DEG considered when the absolute value of log2 fold change > 1 and false discovery rate < 0.05. The list of CEBPA targets was obtained from Harmonizome3.0 website (maintained by the Maayan lab) (34) consisting of the CHEA transcription factor targets.

### Gene Set Enrichment Analysis (GSEA)

GSEA software (35, 36) was performed using default parameters on a pre-ranked list of genes (gene symbols and log2 fold change) with the Hallmark all symbols gene sets, except for the limits of gene set sizes, which were set from 5 to 500.

### Infinium Methylation EPIC arrays

After DNA collection at the indicated time points (two samples per condition), it was shipped to Diagenode Inc. to perform the following downstream steps needed for the Infinium Methylation EPIC array approach. To process the Infinium methylation EPIC array, we followed the workflow from Jovana Maksimovic, Belinda Phipson and Alicia Oshlack as instructed on their vignette (37), until the probe-wise differential methylation analysis, with some modification. Mainly, we used Infinium 850K array annotations (package IlluminaHumanMethylationEPICanno.ilm10b4.hg19) instead of the Infinium 450K in their vignette.

## Results

### AW1-51 induces expression of CEBPA and demethylation of CEBPA promoter in K562 cell line

As previously shown, AW1-51 leads to upregulation of *CEBPA* mRNA 72 hours upon transfection into HepG2 liver cancer cells (26). As *CEBPA* is frequently silenced in various cancers by abnormal DNA methylation of the *locus* (38–40), we studied whether AW1-51-mediated *CEBPA* reactivation could be linked to progressive demethylation of the gene *locus*. To this end, we delivered AW1-51 into the myeloid leukemia cell line K562.

K562 cell line displays low to undetectable levels of *CEBPA* mRNA (21, 41), and lacks CEBPA protein expression due to the presence of the BCR-ABL translocation which prevents translation of *CEBPA* mRNA (42) (Supplementary Figure 1a and 1b). These specific features make K562 cells a suitable model for studying the transcriptional activation induced by AW1-51, without possibly confounding effects linked to *CEBPA* mRNA translation.

As DNA methylation occurs during the S-phase, AW1-51 transfected cells were synchronized at G1/S phase by double thymidine block (25), and *CEBPA* mRNA levels were measured at 24, 48 and 72 hours upon transfection as compared to a negative control (NC) (Figure 1b and 1c). Increased expression of *CEBPA* mRNA at the set timepoints was observed along with parallel changes of the *ecCEBPA* transcript, previously shown to control the *CEBPA locus* DNA methylation (21) (Fig. 1c).

Building on these findings, we sought to investigate whether AW1-51 transcriptional activation would result in a loss of DNA methylation at the target *locus*. *CEBPA* promoter demethylation response was first restated by treating K562 cells with 10 μM 5-Azacitidine (5-Aza) and quantifying *CEBPA* expression 72 hours after treatment. 5-Aza treatment led to an increase of *CEBPA* expression along with a strong demethylation effect in the region between -0.85 to -0.55kb upstream to the transcriptional start site (TSS) (Supplementary Figures 1c and 1d) as shown in previous studies (21), suggesting that the methylation status of this specific region is linked to gene silencing.

Upon transfection with AW1-51, a significant decrease in DNA methylation of the distal promoter located at -0.85 to -0.55kb from the *CEBPA* TSS was detected at 24 and 48 hours after delivery when compared to the non-targeting control (Figure 1d and Supplementary Figure 1e). This effect was lost at 72 hours, suggesting the transient effect of saRNAs, as reported by other studies (26), while no significant changes in 5-hydroxymethylcytosine by (5hmC) by oxidative bisulfite sequencing (OX-BS seq) were observed in the same region (Supplementary Figure 2a and 2b).

Analyses of potential off-target effects by the Infinium Methylation EPIC array platform did not reveal substantial changes of the methylome associated with AW1-51 delivery, indicating a specific on-target effect for the *CEBPA* distal promoter (Supplementary Figure 3a). that is not covered by the Infinium Methylation EPIC array (Supplementary Figure 3b).

### AW1-51 induces demethylation in CEBPA locus and increase of CEBPA mRNA and protein expression in A549

To examine whether the effect of AW1-51 could be extended to cell lines other than K562, we sought for a model wherein the decrease of DNA methylation in the *locus* along with the parallel increase of *CEBPA* mRNA would result in functional expression of the respective protein. *CEBPA* expression is frequently silenced by DNA methylation in lung cancer (14, 43–45), therefore, the lung cancer cell line A549, which is characterized by DNA methylation at the *CEBPA* promoter was used to further investigate the effect of AW1-51 (45). Unlike K562, A549 displays low but detectable levels of *CEBPA* mRNA and protein (14), which can be increased upon treatment with hypomethylating agents (45).

Delivery of AW1-51 into synchronized A549 cells led to an increase of *CEBPA* mRNA at 48- and 72-hour timepoints as compared to the NC saRNA and parallelled by upregulation and stabilization of the *ecCEBPA* transcript (Figure 2a and 2b). Striking changes in cell morphology were noted 120 hours upon AW1-51 delivery (Figure 2c), consistently with those previously reported in lung cancer cell models upon induction of *CEBPA* previously described to be related to cell differentiation (14).

**Figure 2.**
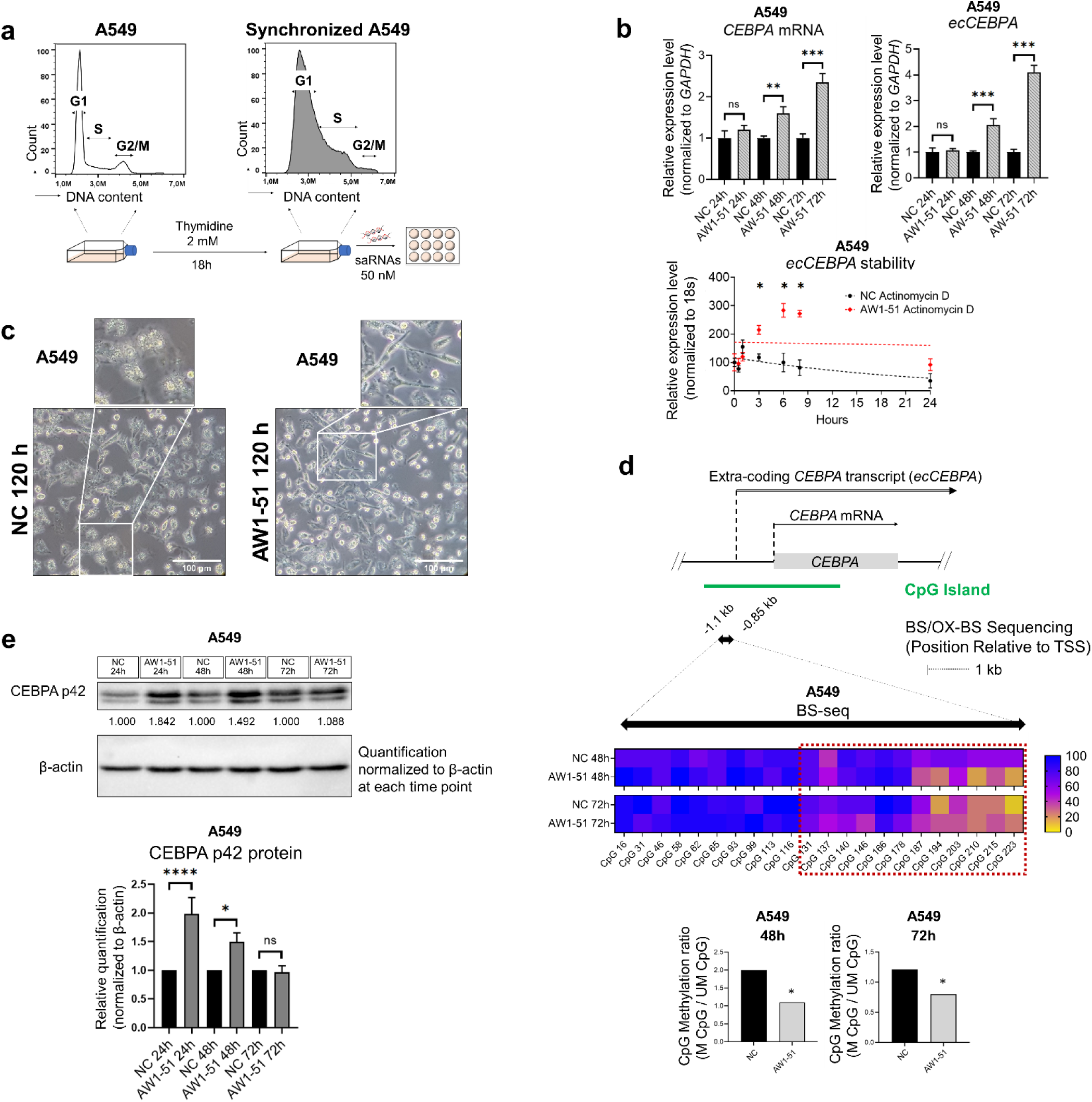
AW1-51 upregulates CEBPA and ecCEBPA transcripts and CEBPA protein under conditions of basal expression. **a** Schematic of saRNA delivery into A549 cells. A549 are seeded 24h prior to thymidine treatment. Only a single block is necessary (2 mM, 1,8h) for G1/S phase arrest. Cells are transfected twice with either AW1-51 or NC: reverse transfection (50 nM), 24h recovery, forward transfection (50 nM). **b,** Higher panel: RT-qPCR results showing CEBPA (left) and ecCEBPA (right) expression from 24h to 72h post transfection in A549 cells (n=3). One-way ANOVA. **: p<O.01, ***: p<0.001, ns: non-significant. Lower panel: ecCEBPA transcript stability in A549 cells treated with 1 µg/mL Actinomycin D assessed by RT-qPCR at 30 min, 1 h, 3 h, 6h, 8h, and 24 h post-treatment. Unpaired t test performed at each time point (n=2). *: p<0.05. **c,** A549 morphological changes with AW1-51 treatment 120h after transfection. 20X magnification, **d,** Region within the CEBPA CpG island (black double-headed arrow, -1.1/-0.85 kb before CEBPA’s TSS) interrogated by BS-seq. Changes in DNA methylation are shown as heatmaps (see detailed BS-seq results in Supplementary Figure 4d). Bar graphs show the ratio between methylated (M) or unmethylated (UM) CpGs to total CpGs within the region included in the dashed line red box. All bisulfite sequenced clones were analyzed by Fisher’s exact test (n=12). *p<0.05. **e,** Higher panel: western blot results showing p42 CEBPA isoform protein expression from 24h to 72h upon the transfection with the AW1-51 or NC. Lower panel: bar graphs represent the relative quantification of CEBPA p42 normalized to β - actin, that was used as our loading control. One-way ANOVA (n= 3). *p<0.05, ****p<0,0001, ns: non-significant.

We then evaluated changes in DNA methylation at the *CEBPA* distal promoter. As the -0.85 to -0.55kb region analyzed in K562 is not methylated in A549, according to combined bisulfite restriction analysis (COBRA) (Supplementary Figure 4a) and ENCODE data, we decided to treat the cells with 5-Aza to identify region(s) methylated in *CEBPA locus* and associated with low *CEBPA* expression. A significant increase of *CEBPA* mRNA levels was measured 96 hours upon treatment with 15μM 5-Aza (Supplementary Figure 4b), while decrease of DNA methylation was detected in a region located between -1.1 and -0.85 kb from the TSS (Supplementary Figure 4c), suggesting that this region plays a role in *CEBPA* expression regulation in this cell line.

We then profiled the DNA methylation across the region spanning -1.1 to -0.85kb from the *CEBPA* TSS at the time points corresponding to *CEBPA* transcripts upregulation following saRNA transfection (48h and 72h) (Figure 2b). After transfection with AW1-51, DNA methylation was significantly reduced compared to the NC both at 48 and 72 hours (Figure 2d and Supplementary Figure 4d). No detectable changes in 5hmC at either time point emerged from OX-BS seq (Supplementary Figure 5a and 5b). Furthermore, genome-wide DNA methylation profiling using the Infinium Methylation EPIC array did not identify additional methylation changes across the A549 genome, confirming that the demethylating effect of AW1-51 was specific to the *CEBPA locus* (Supplementary Figure 6a). However, this sequence did not appear as differentially methylated in the array-based analysis, since the array probes do not overlap with the specific region interrogated by bisulfite sequencing in A549 cells (Supplementary Figure 6b).

Lastly, we assessed the CEBPA protein levels upon AW1-51 transfection at the same time points used to measure the *CEBPA* transcripts levels in A549 cells. While transcripts levels increased gradually (Figure 2b higher panel), we observed an early upregulation of CEBPA protein at 24 hours, followed by a progressive decline from 24 to 72 hours (Figure 2e).

These findings suggest that AW1-51 may also exert effects at the translational level, indicating a potential interplay between saRNAs and early protein expression dynamics.

### A549 exhibited a “CEBPA signature” after transfection with AW1-51

To evaluate the transcriptomic changes and potential off-target effects of AW1-51, we performed RNA sequencing (RNA-seq) 72h after saRNAs transfection in both K562 and A549 cells. Notably, A549 exhibited a significant number of differentially expressed genes (DEGs) unlike K562 (Figure 3a). This difference may be attributed to the restoration of CEBPA protein expression in A549 cells (Figure 2e), which is absent in K562. Indeed, many of the genes upregulated or downregulated in A549 are known *bona-fide* CEBPA targets, whereas no such enrichment was observed in K562, suggesting that AW1-51 activates CEBPA-dependent transcriptional programs, thereby establishing a “CEBPA signature” in A549 cells (Figure 3b, Figure 3c and Supplementary Data 2).

**Figure 3.**
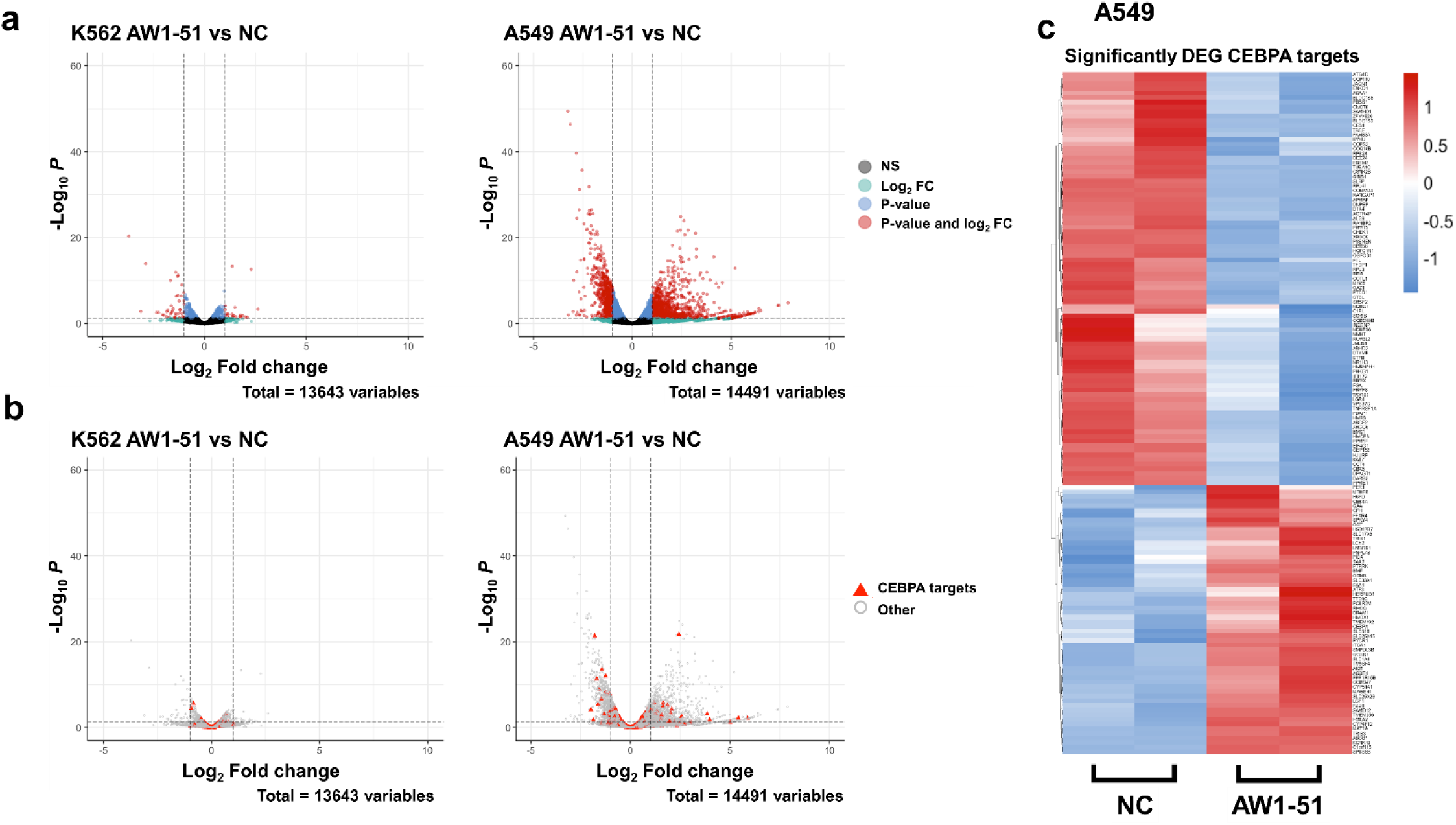
AW1-51 induced a “CEBPA signature” in A549 after transfection. **a,** Volcano plots showing all the differentially expressed genes (DEG) in K562 (left) and A549 (right) when comparing AW1-51 RNA-seq results vs NC RNA-seq results 72h after saRNAs treatment. K562 presented 51 significant DEG and A549 had 4,008 significant DEG. Legend: grey dots: non-significant genes: green dots: genes only satisfying the Log2 fold change threshold parameter (|Log2 fold change| > 1; dashed vertical lines); blue dots: genes only satisfying the p-value parameter (p-value < 0.05; dashed horizontal line); and red dots: significant genes in terms of both Log2 fold change and p-value. **b,** Same volcano plots shown in Figure 3a highlighting CEBPA downstream targets (red triangles). c, Heatmap showing the 147 significant DEGs CEBPA targets when comparing NC and AW1-51.

To further characterize these transcriptomic changes, we performed Gene Set Enrichment Analysis (GSEA) on the significant DEGs in A549. Among the significantly downregulated gene sets, many of them were associated with cell cycle progression and cellular energetics, including mitochondrial metabolism (Supplementary Data 3), both of which are tightly linked to cell proliferation (46). Their downregulation may contribute to the anti-tumor effects of AW1-51 in this context. Regarding the significantly upregulated gene sets, several are enriched for genes involved in the inflammatory response pathway (Supplementary Data 3).

## Discussion

In this study we dissected the mechanism through which AW1-51, a 21 base pairs double stranded saRNA targeting the *CEBPA locus*, elicits transcriptional activation of its target gene through modulation of DNA methylation. Several hypothetical mechanisms of the saRNAs’ activity have been suggested (2–5), yet the precise molecular mode of action remains unclear.

Importantly, AW1-51 is currently under clinical investigation in patients with advanced liver cancer (ClinicalTrials.gov: NCT02716012; EudraCT 2021-005431-23) (8, 9), and in advanced solid tumors (ClinicalTrials.gov: NCT04105335) (10–12), underscoring the translational relevance of these findings and the significance of elucidating the molecular pathways.

We evaluated AW1-51 activity in two cellular models with a distinct *CEBPA* expression profile. In K562, *CEBPA locus* is epigenetically silenced and CEBPA protein suppressed due to BCR-ABL translocation (42), whereas A549 cells express low levels of *CEBPA* mRNA and protein. These contrasting models enabled us to study AW1-51’s effect solely on transcription or on both transcription and translation.

Previous studies have shown that AW1-51, is loaded onto Argonaute 2 (AGO2) and translocated into the nucleus wherein it binds to the *CEBPA locus* via sequence complementarity to increase transcription (26). Here, we report for the first time that AW1-51 also elicits *locus*-specific DNA demethylation at the *CEBPA* promoter in both myeloid leukemia and lung cancer cell lines, with no concurrent changes in 5-hydroxymethylcytosine (5hmC). To our knowledge, this is the first evidence of a saRNA directly influencing DNA methylation at its target *locus* (Figure 4a).

**Figure 4.**
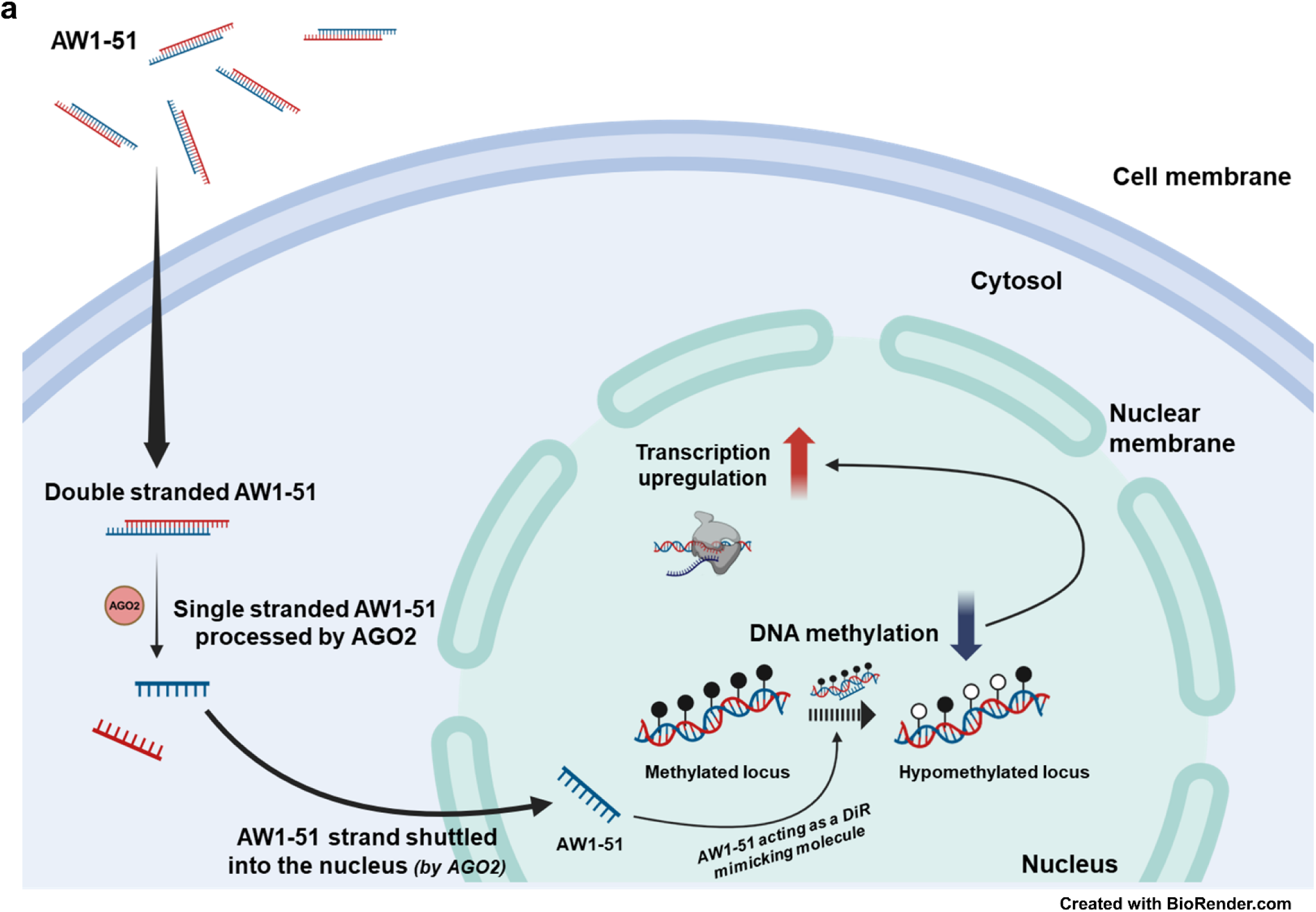
Graphical abstract. Proposed model summarizing the mechanism by which AW1-51 induces *CEBPA* transcriptional upregulation through locus-specific demethylation. Created with BioRender.com

The demethylation effect may be further propagated by upregulation and stabilization of *ecCEBPA*, a DNMT1-interacting RNA (DiR) known to inhibit DNMT1 *in cis* at the *CEBPA locus* (21). Indeed, the increased DNA methylation observed at 72h in K562, paralleled the reduction of *ecCEBPA* levels as compared to earlier time points (24 and 48 hours) upon AW1-51 treatment. These results indicate that AW1-51 may elicit a DiR-mimicking effect by promoting expression of *ecCEBPA*, thereby suggesting an additional mode of action of saRNAs.

Furthermore, AW1-51 induced transient increase of CEBPA protein expression in A549 cell line at early time points, suggesting a possible role in translational regulation. Notably, protein upregulation led to activation of CEBPA downstream pathways in A549, an effect not observed in K562 and consistent with the absence of CEBPA protein. Gene Set Enrichment Analysis (GSEA) of differentially expressed genes in A549 following AW1-51 transfection revealed significant enrichment in pathways associated with cell cycle and cellular energetics (downregulated), and with inflammation (upregulated).

Given the tight link between cell cycle progression and mitochondrial metabolism (47), these data suggest that AW1-51 may exert anti-proliferative effects in cancer cells. This raises the possibility of combining saRNA-based therapy with existing agents targeting inflammatory regulators (48, 49). Interestingly, the early protein upregulation observed in A549 did not mirror the gradual increase in *CEBPA* mRNA, suggesting that AW1-51 may enhance *CEBPA* mRNA translation after transfection. This result hints at an uncharacterized cytosolic role for AW1-51, and other saRNAs whose functions have been largely studied within the nucleus (26, 50, 51). Further investigation is needed to confirm this hypothesis.

The design and the synthetic production of saRNAs capable of upregulating both transcription and translation of disease-relevant genes holds great therapeutic promise. By enabling the reactivation of key regulatory genes implicated in cancer, neurodegenerative, and cardiovascular diseases (52) these molecules pave the way for a novel class of precision therapeutics.

## Data availability statement

The Infinium Methylation EPIC arrays datasets supporting the findings of this work have been submitted to the Gene Expression Omnibus (GEO). These datasets are temporarily private but can be accessed upon reasonable request to the corresponding author. The GEO accession number will be made publicly available upon journal publication.

The RNA-seq datasets supporting the findings of this work have been submitted to the Gene Expression Omnibus (GEO). These datasets are temporarily private but can be accessed upon reasonable request to the corresponding author. The GEO accession number will be made publicly available upon journal publication.

## Acknowledgments and funding

This work has been supported by the ADR lab funding: the ACS RSG-23-1036643-01-DMC, the NIH/NIDDK R01DK136116, the HIRM Pilot Award 2021, NCI R00 CA188595, Italian Association for Cancer Research (AIRC) Start-Up grant #2014-15347. JRPM was supported by an Alfonso Martin Escudero Foundation Postdoctoral Fellowship, an EMBO Scientific Exchange Grant (9947) and a Predoctoral FPU Fellowship from the Spanish Ministry of Universities (FPU18/03709). SU was supported by a National Institutes of Health, National Research Service Award (NRSA) [5T32HL007917-24]. DGT was supported by 1P01HL131477-6 A1 (NIH/NHLBI) and 2023 Grant # 2023_TJF_02 from the AGA/Jenzabar Foundation. We would like to thank the BIDMC core facilities for their support in this work.

## Author contributions

ADR conceived the study, coordinated the scientific team, and allocated the funding for the project; GG, JRP-M, MB and ML generated most of the experimental data; ADR, GG, JRP-M, MB, ML, LR, SU, DD, AKE designed experiments, analyzed data, and wrote the manuscript; LR and MAB performed bioinformatics analyses; NAH provided the AW1-51 strands sequences; ADR, GM, SSK, DGT, BG, MPP, NAH and ADB provided scientific advice and reviewed the manuscript. All authors contributed with suggestions after a critical reading of the draft and approve its submission for publication.

## Conflict of interest statement

ADR is a founder and scientific advisor of Aptadir Therapeutics; all interests are outside the submitted work. DGT is a founder and scientific advisor of Aptadir Therapeutics; all interests are outside the submitted work. NAH is a founder and shareholder of MiNA Therapeutics Limited and currently receives no financial support from MiNA Therapeutics by way of personal payments, grants, or expenses. SSK reports receiving consulting fees and research support from Taiho Therapeutics; research support from Boehringer Ingelheim, MiRXES, and Johnson & Johnson; honoraria from AstraZeneca, Boehringer Ingelheim, Bristol Myers Squibb, Chugai Pharmaceutical, and Takeda Pharmaceuticals; royalties from Signosis and Life Technologies; all interests are outside the submitted work.

