## Supplementary Figures 1-6 for "Small activating RNA AW1-51 (CEBPA-51) elicits targeted DNA demethylation to promote gene activation"

Supplementary Figure 1

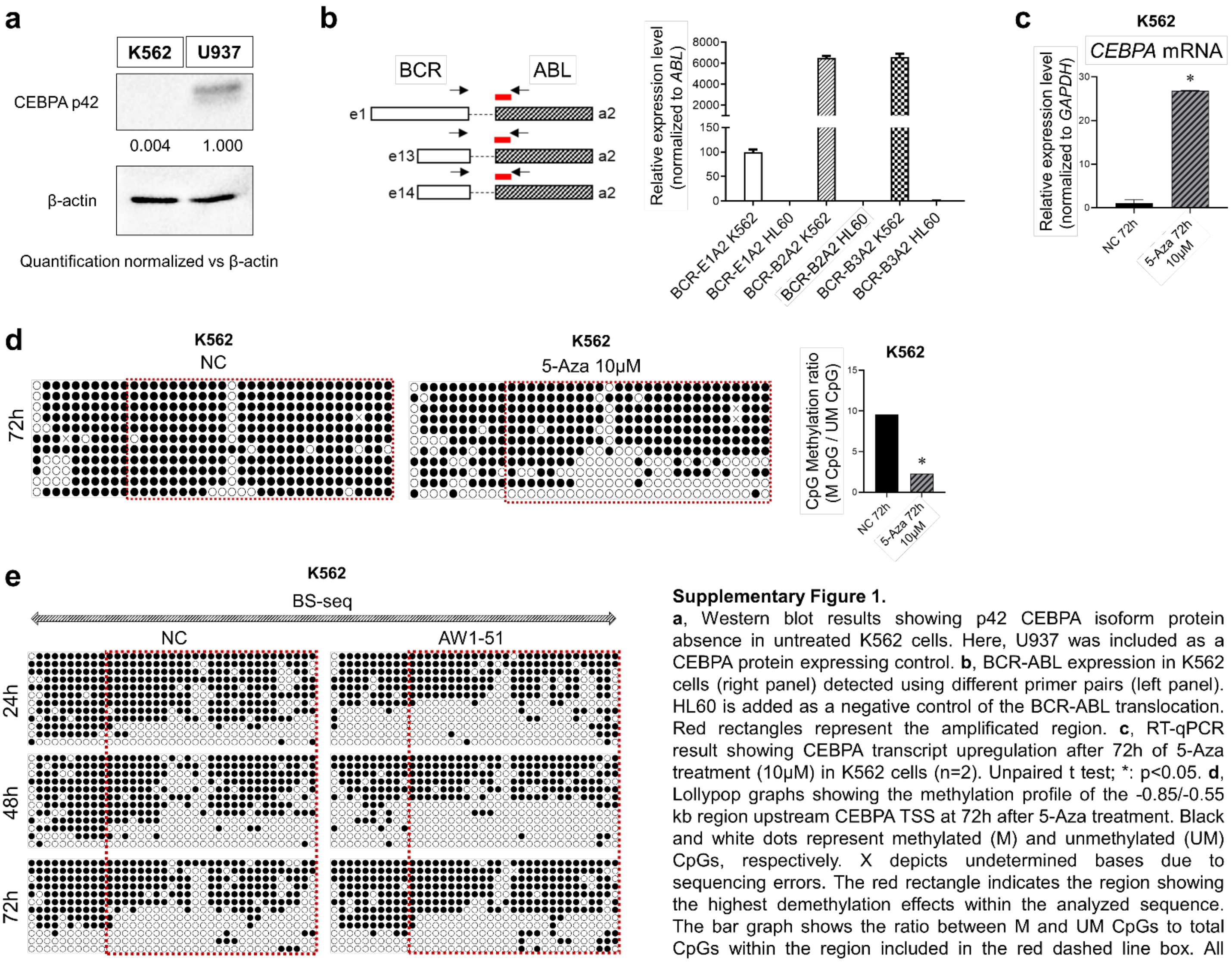


**Supplementary Figure 1.**

**a**, Western blot results showing p42 CEBPA isoform protein absence in untreated K562 cells. Here, U937 was included as a CEBPA protein expressing control. **b**, BCR-ABL expression in K562 cells (right panel) detected using different primer pairs (left panel). HL60 is added as a negative control of the BCR-ABL translocation. Red rectangles represent the amplificated region. **c**, RT-qPCR result showing CEBPA transcript upregulation after 72h of 5-Aza treatment (10µM) in K562 cells (n=2). Unpaired t test; *: p<0.05. **d**, Lollypop graphs showing the methylation profile of the -0.85/-0.55 kb region upstream CEBPA TSS at 72h after 5-Aza treatment. Black and white dots represent methylated (M) and unmethylated (UM) CpGs, respectively. X depicts undetermined bases due to sequencing errors. The red rectangle indicates the region showing the highest demethylation effects within the analyzed sequence. The bar graph shows the ratio between M and UM CpGs to total CpGs within the region included in the red dashed line box. All bisulfite sequenced clones were analyzed by Fisher's exact test (n=12); *: p<0.05. **e**, Lollypop graphs showing the detailed methylation (BS-seq) profile of -0.85/-0.55 kb region upstream CEBPA TSS after NC or AW1-51 transfection at the indicated timepoints. Black and white dots represent methylated (M) and unmethylated (UM) CpGs, respectively. X depicts undetermined bases due to sequencing errors. The red rectangle indicates the region showing the highest demethylation effects within the analyzed sequence. All bisulfite sequenced clones were analyzed by Fisher's exact test (n=12); *: p<0.05.

### Supplementary Figure 2

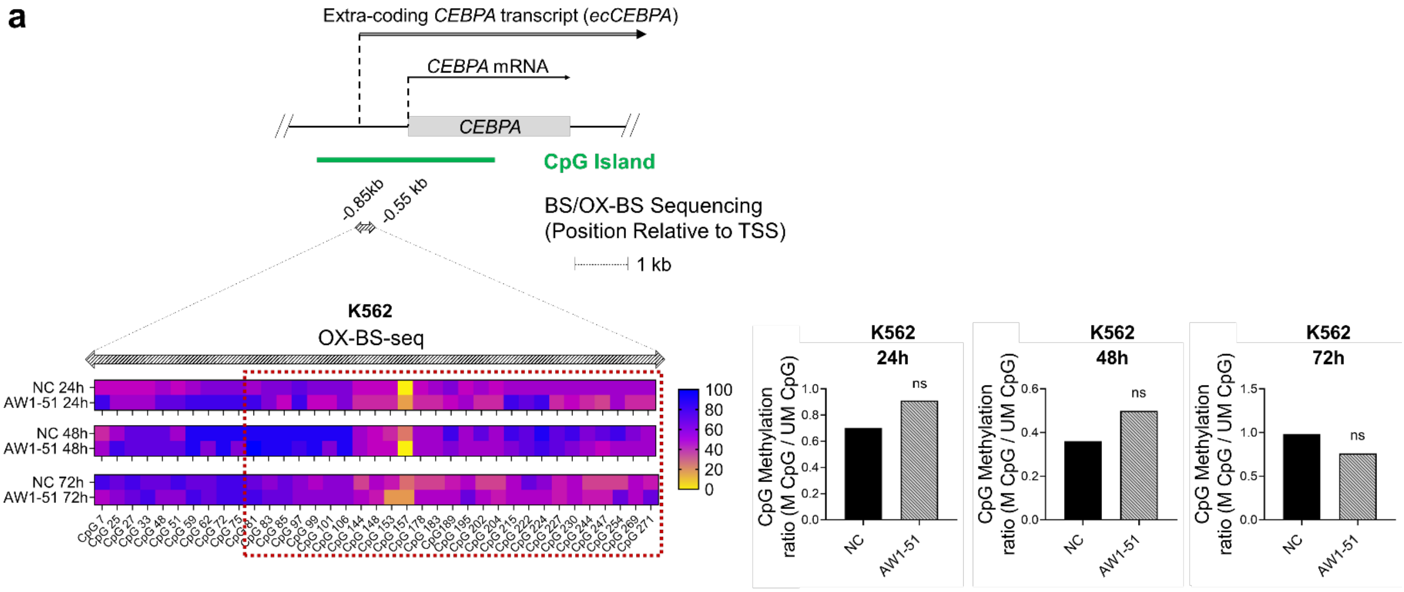


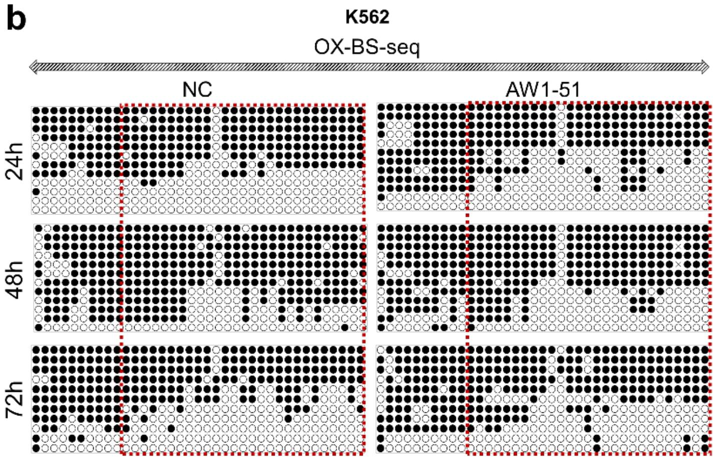


**Supplementary Figure 2.**
**a**, Region within the CEBPA CpG island (striped double-headed arrow, -0.85/-0.55kb before CEBPA's TSS) interrogated by oxidative bisulfite sequencing (OX-BS-seq) in K562 cell. Changes in DNA hydroxymethylation are shown as heatmaps. Bar graphs show the ratio between hydroxymethylated (hM) and non-hydroxymethylated (NhM) CpGs to total CpGs within the region included in the red dashed line box. All bisulfite sequenced clones were analyzed by Fisher's exact test (n=12); ns: non-significant. **b**, Lollypop graphs showing the detailed DNA hydroxymethylation (OX-BS-seq) profile of the -0.85/-0.55 kb region upstream CEBPA TSS in K562 after NC or AW1-51 transfection at the indicated timepoints. Black and white dots represent hydroxymethylated (hM) and non-hydroxymethylated (NhM) CpGs, respectively. X depicts undetermined bases due to sequencing errors. The red rectangle shows the region showing the highest demethylation effects within the analyzed sequence (BS-seq, same region than red dashed line box in Figure 1d, Supplementary Figure 1c and 1d).

Supplementary Figure 3

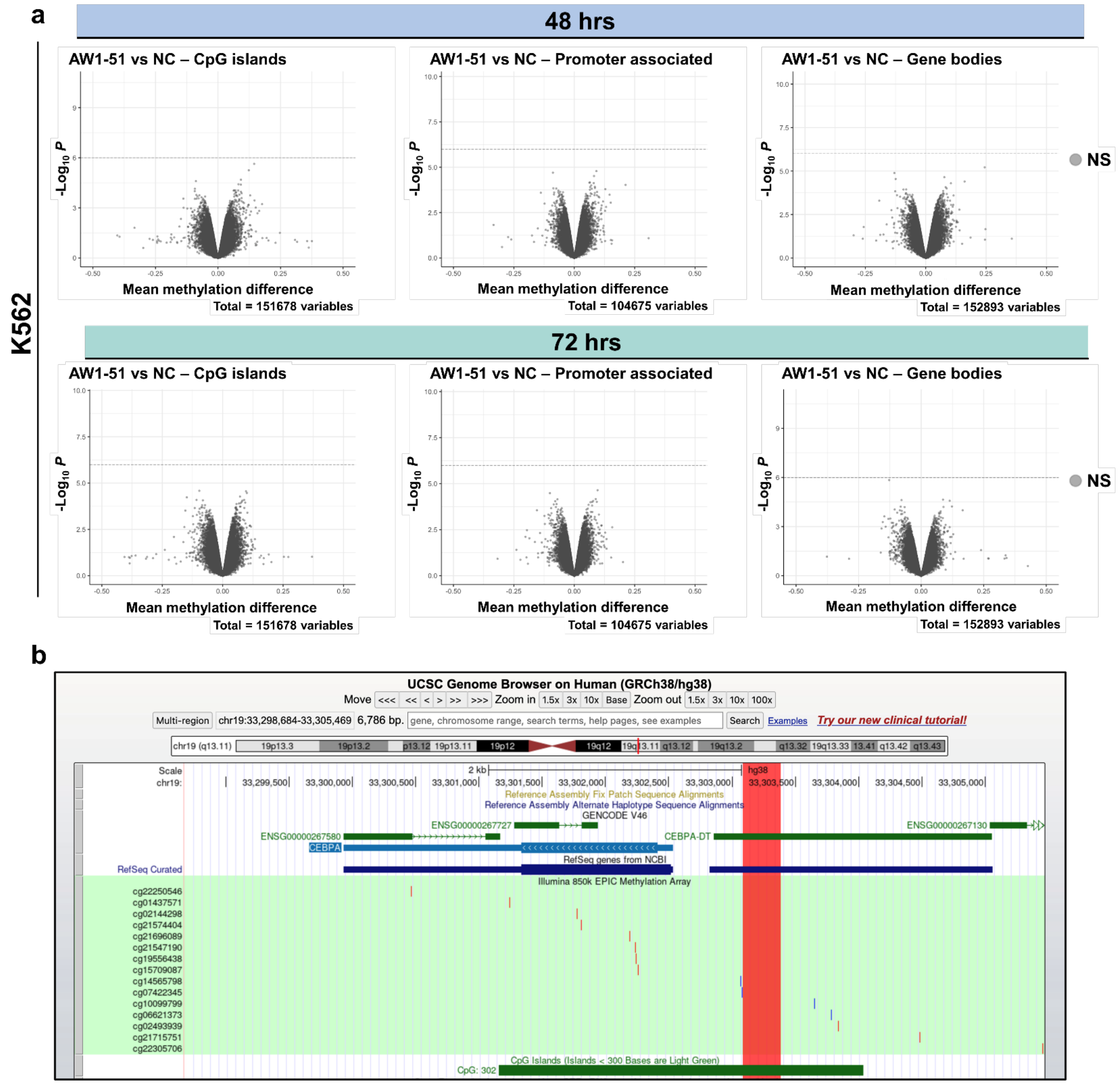


**Supplementary Figure 3.**
**a**, Infinium Methylation EPIC array analysis results showing the differential methylation profile in AW1-51 vs NC in K562 at 48h (top) and 72h (bottom) post transfection in CpG Islands, promoter regions and genes bodies, as indicated. None of the loci analyzed by this array were significant (horizontal dashed line). **b**, Schematic genome representation showing the Infinium Methylation EPIC array probes (highlighted with a light green background) that match around the CEBPA locus. The vertical red rectangle is highlighting the region that was assessed by BS-seq and OX-BS-seq in K562 (-0.85 to -0.55kb from CEBPA's TSS). None of the probes anneal within this specific region. Genome assembly: GRCh38/hg38. Screenshot obtained from UCSC Genome Browser.

### Supplementary Figure 4

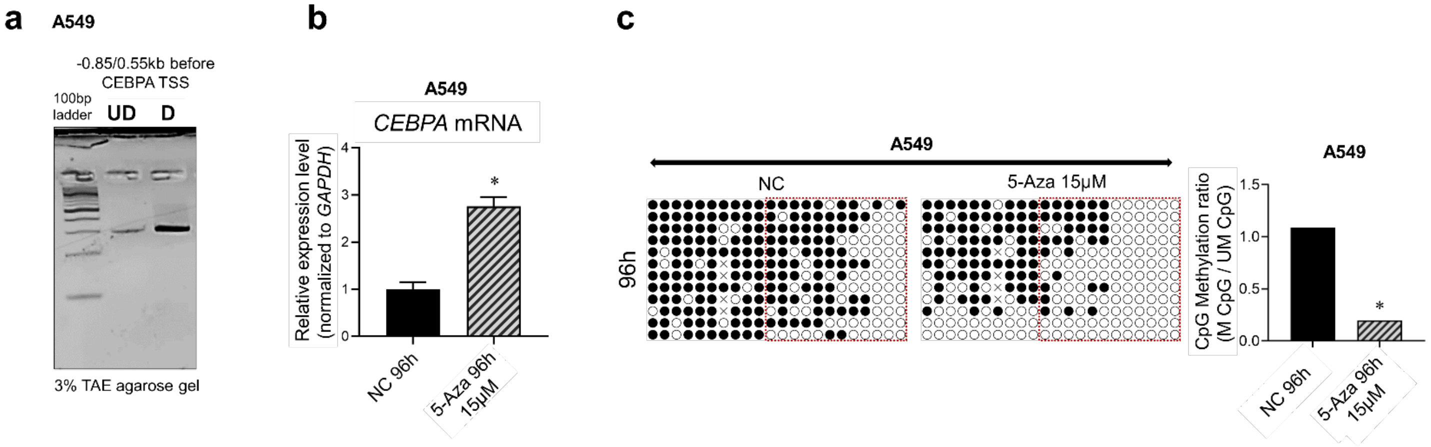


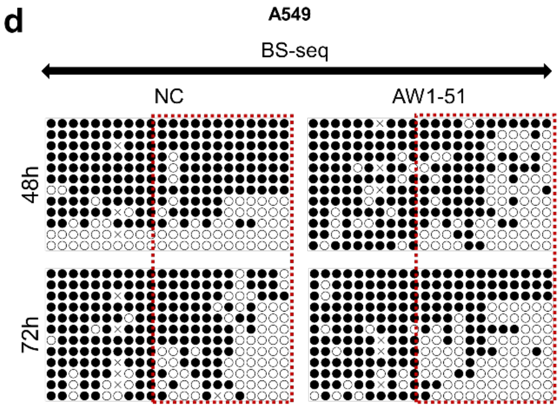


**Supplementary Figure 4.**

**a**, DNA methylation profile analysis by combined bisulfite restriction analysis (COBRA) in the -0.85/-0.55 kb region upstream CEBPA TSS. UD and D represent the undigested and BstUI-digested product, respectively. **b**, RT-qPCR result showing CEBPA transcript upregulation after 96h of 5-Aza treatment (15µM) in A549 cells (n=2). Unpaired t test; *: p<0.05. **c**, Lollypop graphs showing the methylation profile of the -1.1/-0.85 kb region upstream CEBPA TSS 96h after 5-Aza treatment. Black and white dots represent methylated (M) and unmethylated (UM) CpGs, respectively. X depicts undetermined bases due to sequencing errors. The red rectangle indicates the region showing the highest demethylation effects within the analyzed sequence. The bar graph shows the ratio between M and UM CpGs to total CpGs within the region included in the dashed line red box. All bisulfite sequenced clones were analyzed by Fisher's exact test (n=12); *: p<0.05. **d**, Lollypop graphs showing the methylation (BS-seq) profile of the -1.1/-0.85 kb region upstream CEBPA TSS after NC or AW1-51 transfection at the indicated timepoints. Black and white dots represent methylated (M) and unmethylated (UM) CpGs, respectively. X depicts undetermined bases due to sequencing errors. The red rectangle indicates the region showing the highest demethylation effects within the analyzed sequence. All bisulfite sequenced clones were analyzed by Fisher's exact test (n=12); *: p<0.05.

Supplementary Figure 5

**a**

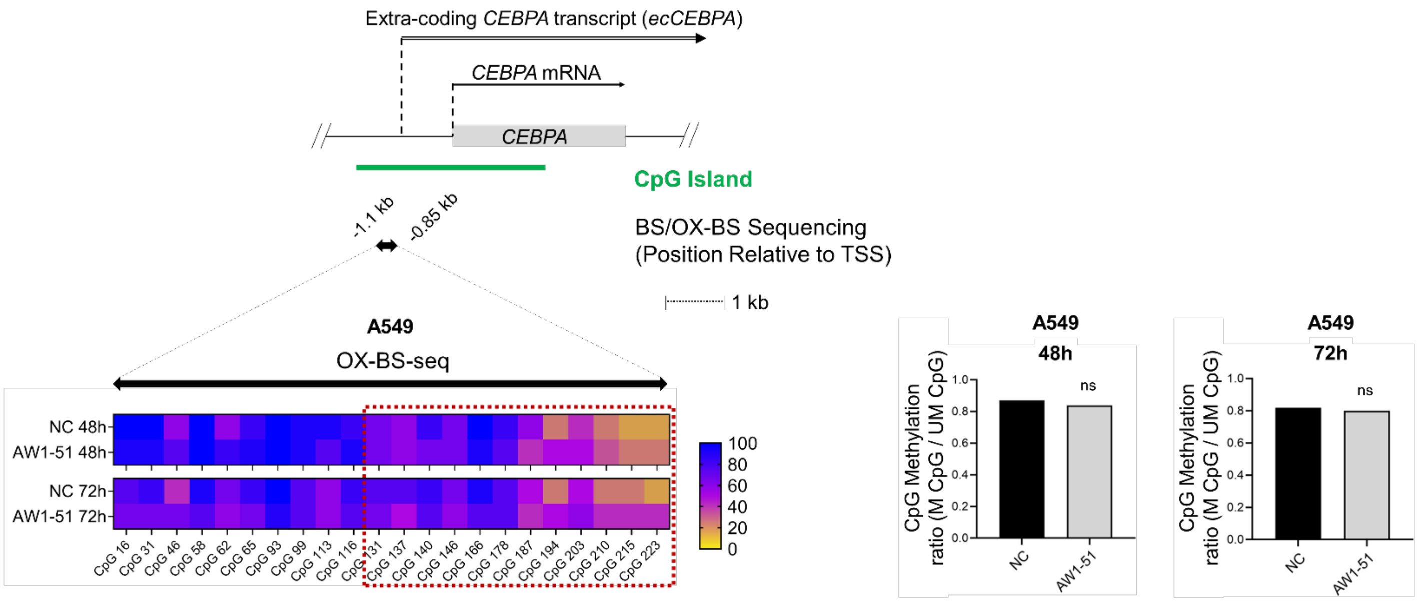


**b**

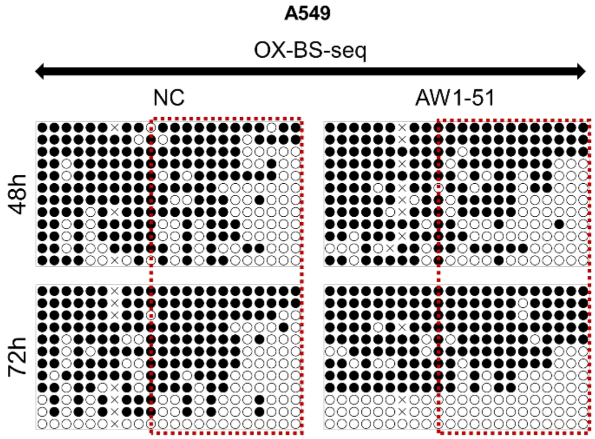


**Supplementary Figure 5.**
**a**, Region within the CEBPA CpG island (black double-headed arrow, -1.1/-0.85 kb before CEBPA's TSS) interrogated by OX-BS-seq in A549 cell. Changes in DNA hydroxymethylation are shown as heatmaps. Bar graphs show the ratio between hydroxymethylated (hM) and non-hydroxymethylated (NhM) CpGs to total CpGs within the region included in the red dashed line box. All bisulfite sequenced clones were analyzed by Fisher's exact test (n=12); ns: non-significant. **b**, Lollypop graphs showing the detailed DNA hydroxymethylation (OX-BS-seq) profile of the -1.1/-0.85 kb region upstream CEBPA TSS in A549 after NC or AW1-51 transfection at the indicated timepoints. Black and white dots represent hydroxymethylated (hM) and non-hydroxymethylated (NhM) CpGs, respectively. X depicts undetermined bases due to sequencing errors. The red rectangle shows the region showing the highest demethylation effects within the analyzed sequence (BS-seq, same region than red dashed line box in Figure 2d, Supplementary Figure 4c and 4d).

Supplementary Figure 6

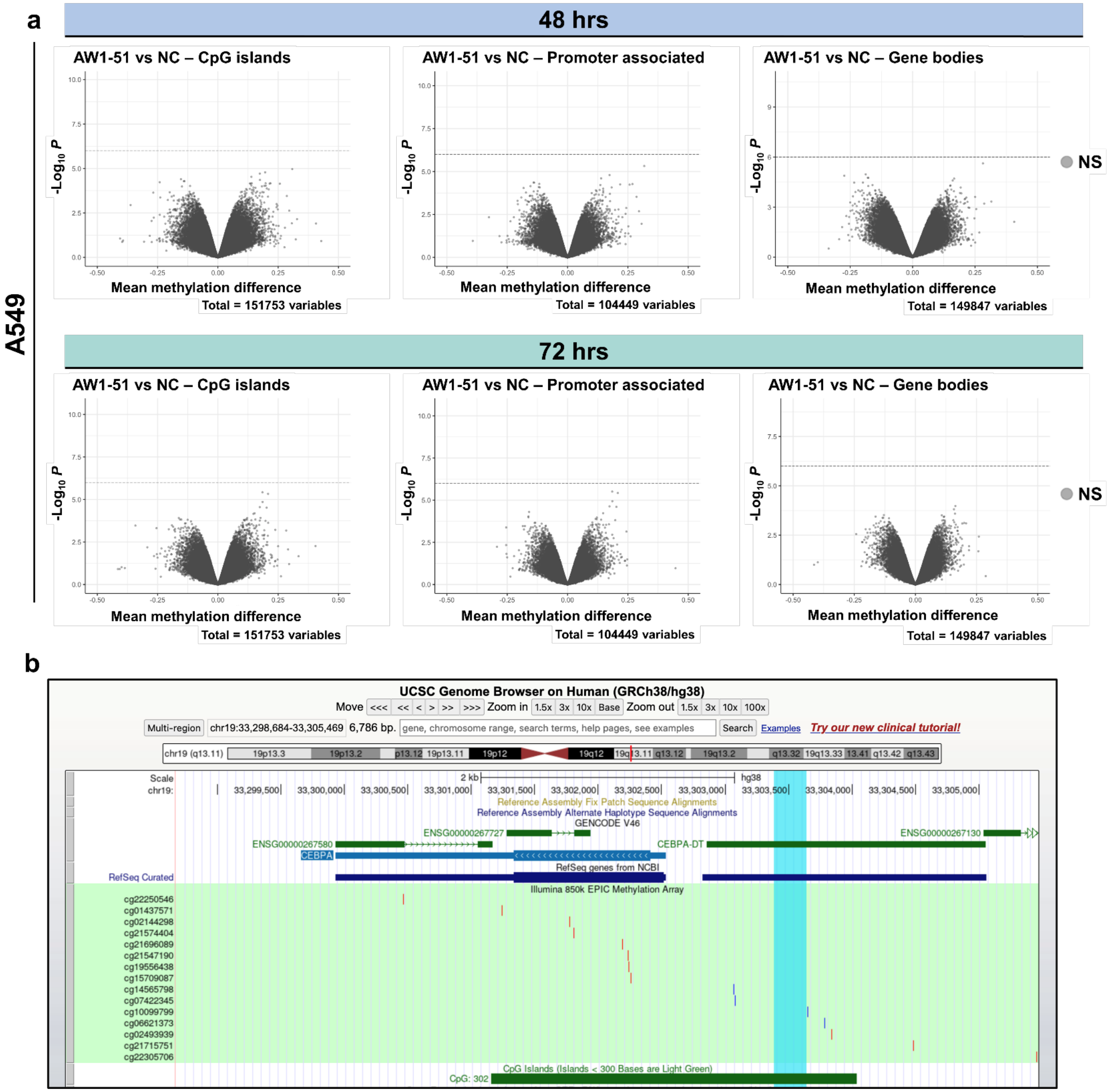


**Supplementary Figure 6.**

**a**, Infinium Methylation EPIC array analysis results showing the differential methylation profile in AW1-51 vs NC in A549 at 48h (top) and 72h (bottom) post transfection in CpG Islands, promoter regions, and genes bodies, as indicated. None of the loci analyzed by this array were significant (horizontal dashed line). **b**, Schematic genome representation showing the Infinium Methylation EPIC array probes (highlighted with a light green background) that match around the CEBPA locus. The vertical cyan rectangle is highlighting the region that was assessed by BS-seq and OX-BS-seq in A549 (-1.1 to -0.85kb from CEBPA's TSS). None of the probes anneal within this specific region. Genome assembly: GRCh38/hg38. Screenshot obtained from UCSC Genome Browser.
