## Supplementary Data 1 for "Small activating RNA AW1-51 (CEBPA-51) elicits targeted DNA demethylation to promote gene activation"

***Small activating RNAs (saRNAs) sequences***

*AW1-51*

Sense strand: 5’ GACCAGUGACAAUGACCGCUU 3’

Antisense strand: 5’ GCGGUCAUUGUCACUGGUCUU 3’

*Negative control (NC) saRNA*

Sense strand: 5’ UCGAAGUACUCAGCGUAAGUU 3’

Antisense strand: 5’ CUUACGCUGAGUACUUCGAUU 3’

***Bisulfite sequencing, oxidative bisulfite sequencing and combined bisulfite restriction analysis (COBRA) primers:***

BS-seq region -0.85 to -0.55kb from the TSS of CEBPA:

FW CEBPA_0.85kb: 5’ TAGTTTYGTTAGTTTGGGGGGTTT 3’

RV CEBPA_0.85kb:5’ TCTAATCTCCAAACTACCCCTATA 3’

BS-seq region -1.1 to 0.85kb from the TSS of *CEBPA*:

FW CEBPA_1.1kb: 5’ TATTTAAGGGGTTTTAGG 3’

RV CEBPA_1.1kb: 5’ AAAAACAAACTTAACTCTAA 3’

***RT-qPCR primers and TaqMan probes sequences:***

*CEBPA*

FW CEBPA: 5’ TCGGTGGACAAGAACAG 3’

RV CEBPA: 5’ GCAGGCGGTCATTG 3’

TaqMan Probe CEBPA: 5’ ACAAGGCCAAGCAGCGC 3’; modifications: 5’ FAM / 3’ BHQ-1.

*ecCEBPA*

FW ecCEBPA: 5’ GGTTGTCTGTGGGCCAGGTCA 3’

RV ecCEBPA: 5’ AGAGCTCATGAAAGTCAGGATTG 3’

TaqMan Probe: 5’ AATAATACAGCATTTTCCCTGGCGG-3’

*GAPDH*

We used “Human GAPD (GAPDH) Endogenous Control (VIC™/TAMRA™ probe, primer limited)”, Applied Biosystems.

*18s*

We used “Eukaryotic 18S rRNA Endogenous Control (VIC™/TAMRA™ probe, primer limited)”, Applied Biosystems.
